# Monitoring Resilience in Bursts

**DOI:** 10.1101/2023.10.10.561665

**Authors:** Clara Delecroix, Egbert H van Nes, Marten Scheffer, Ingrid A van de Leemput

## Abstract

The possibility to anticipate critical transitions through detecting loss of resilience has attracted attention in a variety of fields. Resilience indicators rely on the mathematical concept of critical slowing down, which means that a system recovers increasingly slowly from external perturbations when approaching a tipping point. This decrease in recovery rate can be reflected in rising autocorrelation and variance in data. To test whether resilience is changing, resilience indicators are often calculated using a moving window in long, continuous time series of the system. However, for some systems it may be more feasible to collect several high-resolution time series in short periods of time, i.e. in bursts. Resilience indicators can then be calculated to detect a change of resilience in a system between such bursts. Here, we compare the performance of both methods using simulated data, and showcase possible use of bursts in a case-study using mood data to anticipate depression in a patient. Using the same number of data points, the burst approach outperformed the moving window method, suggesting that it is possible to down-sample the continuous time series and still signal of an upcoming transition. We suggest guidelines to design an optimal sampling strategy. Our results imply that using bursts of data instead of continuous time series may improve the capacity to detect changes in systems’ resilience. This method is promising for a variety of fields, such as human health, epidemiology, or ecology, where continuous monitoring is costly or unfeasible.

**Significance statement:** Gauging the risk of tipping points is of great relevance in complex systems ranging from health to climate, and ecosystems. For this purpose, dynamical indicators of resilience are being derived from long continuous time series to monitor the system and obtain early warning signals. However, gathering such data is often prohibitively expensive or practically unfeasible. Here we show that collecting data in brief, intense bursts may often solve the problem, making it possible to estimate change in resilience between the bursts with relatively high precision. This may be particularly useful for monitoring resilience of humans or animals, where brief time series of blood pressure, balance, mood or other relevant markers may be collected relatively easily to help estimating systemic resilience.

## Introduction

Critical transitions happen when a self-reinforcing feedback pushes a runaway change to a contrasting state once a critical threshold is passed. Such changes can be induced by a gradual change in an underlying condition, bringing the system closer to a critical threshold (1). A range of complex systems have been suggested to undergo critical transitions. Examples include the start and end of depression episodes (2, 3), extinction of populations in ecosystems (4) and disease outbreaks (5).

Systems that gradually approach a critical threshold may display signs of critical slowing down: they recover from external perturbations increasingly slowly. This is related to a decrease in their resilience, defined as the magnitude of a perturbation that a system can stand without tipping to an alternative attractor (6). Thus, the recovery rate of a system pushed by an external perturbation indicates the system’s distance to the critical threshold (7, 8). As it is often impossible to measure the recovery rate directly (e.g., by perturbing the system experimentally), statistical metrics that capture changes in the fluctuations resulting from the natural regime of stochastic perturbations are used as a proxy. Autocorrelation and variance are the most commonly used indicators of critical slowing down in such data. Both indicators are expected to increase when a system gradually approaches a critical transition (9, 10).

In long time series of a system’s representative variable, a moving window is typically used to calculate trends in resilience indicators (10). In other words, indicators are calculated repeatedly in overlapping subsets of the data to reveal their evolution over time. For example, Dakos et al. (2008) demonstrated that resilience indicators calculated in climate time series showed a significant trend before abrupt changes. Harris, Hay and Drake (2020) detected changes in resilience indicators in incidence time series prior to the re-emergence of malaria in Kericho, Kenya, and van de Leemput et al. (2014) and Wichers and Groot (2016) detected critical slowing down prior to the start of depressive episodes.

Although resilience indicators are a valuable model-free anticipation method, they are not suitable for all situations. (9) listed various situations where critical slowing down cannot be expected prior to an abrupt shift. Most importantly, the moving window method of resilience indicators cannot be expected to serve as a warning signal if there is no gradual change in an underlying condition. Even if a gradual change in resilience exists, long high-resolution equidistant time series with small observation errors are required for the method to work. The number of data points should be high enough to observe a change in the system (10). Additionally, for measuring autocorrelation, the resolution of the time series should be high compared to the time scale of the system’s dynamics (9). As monitoring is generally costly, such data requirements are rarely met

for some systems. When patients are involved, collecting long, uninterrupted time series may also be challenging. For instance, only in exceptional cases a time series of the mood of a patient is long enough to detect a change in resilience indicators prior to depression episodes (3). The required data length and resolution (e.g., up to ten times a day for 239 days (3)) is very demanding and will lead to high dropout rates.

Here, we explore the alternative to use bursts of data instead of continuous time series to monitor a potential loss of resilience in a system. We define bursts as short periods of intense monitoring in a system, leading to multiple short time series of high resolution. We estimate resilience within bursts using autocorrelation and variance and subsequently compare bursts. We discuss when monitoring in bursts can be more efficient, but choices remain on the number and duration of the bursts, their resolution, and the interval between two bursts. In some cases, the per-sample costs can be lower than the per-burst costs, for instance, if an expensive automatic device needs to be reserved for the sampling duration. In contrast, in other cases, the total monitoring costs are solely determined by the costs of the individual samples. As the costs per burst and sample vary and can influence the way to allocate sampling resources, we investigate the effect of various scenarios on the performance of the burst approach for monitoring resilience.

We used a well-studied model with a tipping point to generate high-resolution time series. From those time series, we extract subsamples to systematically compare different approaches for monitoring resilience. For a complementary empirical example, we apply the burst approach to a long time series of self-reported moods used to predict the onset of a depressive episode (3). Finally, we formulate advice on how to monitor resilience depending on the system setup and available monitoring resources.

## Methods

### Synthetic data

To compare the performance of resilience indicators with both the rolling window and burst approach, we first generated long-term high-resolution time series using a commonly used model with critical transitions, referred to as the overgrazing model.

We used a simple one-variable model described by May et al. (13). The model is a generic stochastic differential equation model capturing the growth dynamics of vegetation under grazing pressure (equation 1).

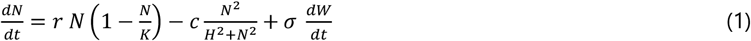

The vegetation *N* grows logistically, and the grazing rate follows a sigmoidal saturation curve. At low vegetation density, grazing is limited by vegetation availability, while at high vegetation density, grazing is limited by the digestion rate of herbivores. In other words, the grazing rate increases with vegetation density until it reaches a maximum. *N* represents the density of vegetation, parameter r the maximum growth rate, *K* the carrying capacity of the population growth, *c* the maximum grazing rate and *H* the grazing saturation constant.

Stochasticity is included in the model to reproduce natural fluctuations as additive noise. We used a Wiener process (dW/dt) with a fixed standard deviation *σ*. This model was implemented in Grind for Matlab (14).

For the default parameter settings used in this paper, the deterministic version of the model (σ=0) displays two-fold bifurcations (F1 and F2 in Figure S1). The high-density vegetation state undergoes a critical transition when the maximum grazing rate c crosses the critical threshold F1.

To simulate a loss of resilience of the high vegetation state, we generated long time series with *c* gradually increasing towards its critical threshold, from 0.5 to 2.2. Other parameter values are fixed on *r*=1, *K* = 10, *H*=1, and *σ*=0.5. The model was simulated for 8000 days with three data points per day, resulting in time series with 24000 data points. A number of 100 stochastic realizations of the time series were simulated to quantify the performance over 100 repetitions. As a control, we generated time series with a fixed *c*=0.5, representing a system with steady resilience.

After simulation, we simulated measurement error by adding a random number from a standard normal distribution (mean 0 and standard deviation σME 0.5) to each data point in the generated time series.

### Sub-sampling the master time series

We see the generated high-resolution time series as representing the actual dynamics of a system and use the resulting ‘master’ time series to sample 1) bursts for the bursts approach and 2) continuous time series for the rolling window approach by using subsets of the data.

To make a fair comparison between approaches, we tried to keep the total number of samples the same for each method (Table 1). When comparing the burst approach with the rolling window approach, this implies that in the rolling window approach, either the master time series is truncated (scenario I) or the data resolution is lower (scenario II). The former leads to a truncated control parameter range (Δc) over the time series. To be complete, we also did the rolling window analysis for the whole master time series (scenario III), although this implies that we use more samples in this analysis. Thus, for each scenario, two of the three sampling choices (Δ1, Δc and L) were the same for both approaches and one differed (Table 1).

**Table 1.** Summary of values for the number of data points (L), the data resolution (Δ1) and the difference in resilience (Δc) for the burst approach and the three scenarios of the rolling window approach.

| | $\Delta_1$ (sampling interval) | $\Delta_c$ (gradient of c values) | L (total number of data points) |
| --- | --- | --- | --- |
| Burst approach | 1 day | 1.7 | 1000 |
| Rolling window approach<br>Scenario I, truncating | 1 day | 0.21 | 1000 |
| Rolling window approach<br>Scenario II, thinning | 24 days | 1.7 | 1000 |
| Rolling window approach<br>Scenario III, gold-standard | 1 day | 1.7 | 8000 |

We also investigated the effect of several sampling choices on the performance of the burst method (Figure 1): the interval between two samples in a burst Δ1 (further referred to as sampling interval) and the interval between two bursts Δ2, which is proportional to the difference in resilience (i.e., because we assume an underlying linear loss of resilience over time), quantified as the difference in bifurcation parameter between the bursts Δc. Additionally, we varied the number of bursts *n*, the number of samples in a burst *λ*, and the total number of samples collected *L=n λ*.

**Figure 1.**
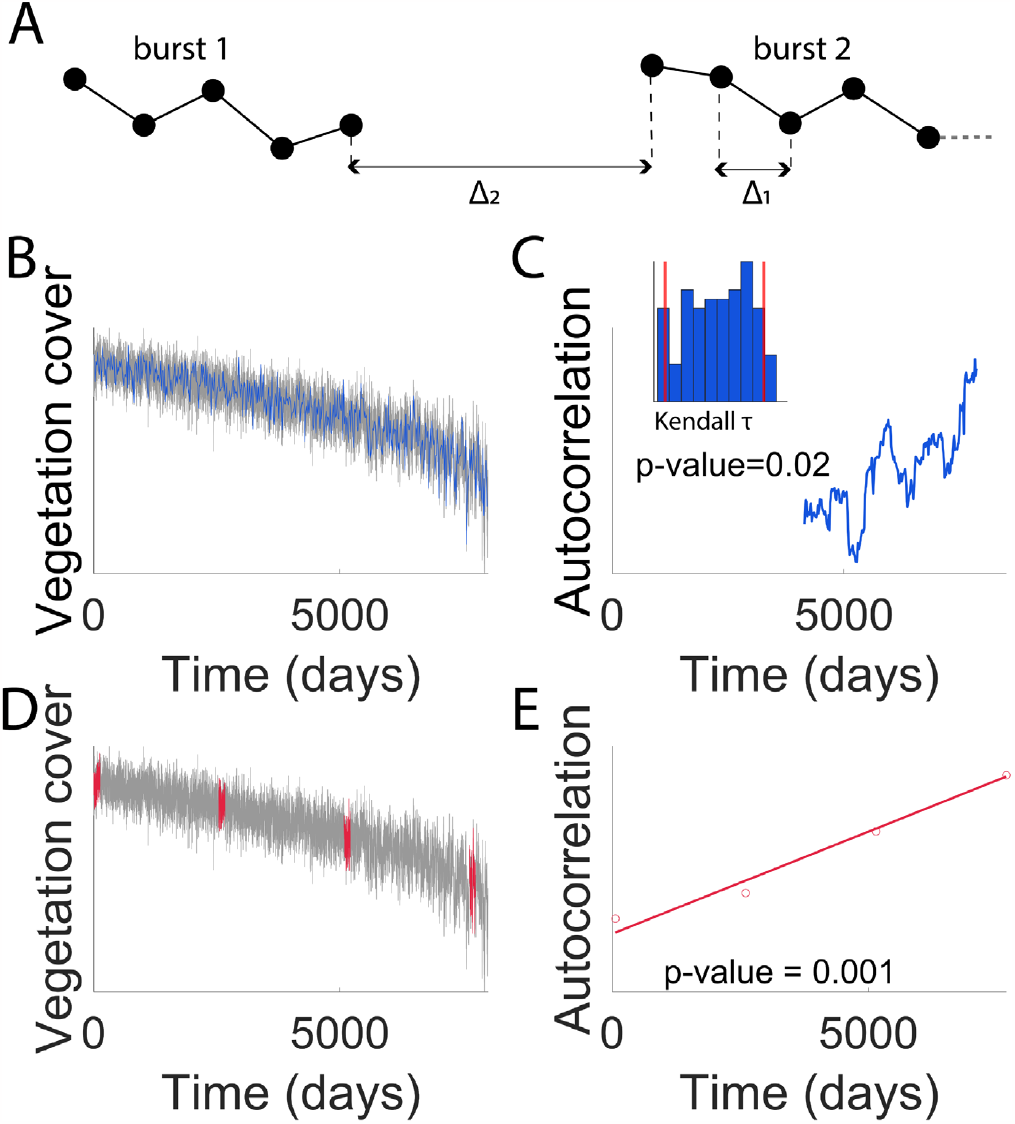
Summary of the types of monitoring to anticipate a transition, with associated analysis methods. (A) Graphical representation of monitoring parameters of interest, Δ1 and Δ2. The circles symbolize the collected samples. (B) Example of samples collected continuously for the rolling window approach. Master time series in grey and sampled time series in blue. (C) Measure of the autocorrelation using a rolling window for the dataset represented in B. The histogram represents the distribution of Kendall tau values in the datasets of the null model. (D) Example of samples collected in 4 bursts for the burst approach. Master time series in grey and sampled bursts in red. (E) Graphical representation of the method used to detect a loss of resilience between several bursts for the dataset represented in D. For each burst, we estimate the indicator. Then, we estimate the slope of the regression for the set of points. We also calculate the p-value of the slope being different from zero.

### Empirical data

In addition to the simulation study, we used a real dataset to compare the proposed burst approach to the conventional method (3, 15). In this study, a patient subject to depressive episodes monitored his daily life experiences up to ten times a day at a random moment for 239 days through a 50-item questionnaire. On average, the interval between two data points was 3 hours and 40 minutes, and 6.2 data points were collected daily. The study comprised five phases: a baseline period of 28 days, a double-blind period with no reduction of the antidepressant intake, a period during which the patient’s antidepressant intake was progressively reduced in double-blind, a post-assessment period, and a follow-up period. The monitoring times were randomly chosen daily, thus, the data points are not equidistant. Additionally, the patient’s depressive symptoms were measured on Mondays, and a significant shift in depressive symptoms was detected on day 127 of the experiments. At the end, 829 data points are available before the shift.

In this study, we focused on the items pertaining to mood states, self-esteem, and physical condition. More specifically, we focused on negative feelings (*feeling down, irritated, lonely, anxious, suspicious, guilty, doubtful, restless, agitated, worried, ashamed, doubting myself, tired, in pain, dizzy, having a dry mouth*, and *nauseous*) as previous work suggests that changing fluctuations in those symptoms may best signal an upcoming depressive episode (2). These affects were all correlated positively with one another. We discarded the items *dizzy, dry mouth*, and *nauseous* as no variation was shown in these affects. With the remaining 14 items, we performed a PCA as a data reduction technique to obtain one time series of 829 data points before the shift. The PCA coefficients were calculated on the baseline period only (the first 28 days, 177 data points), and the time series were projected in the direction of the first principal component. Similarly to the simulation study, we subsampled from the master time series to compare the burst and the rolling window approach. We used the previously defined down-sampling scenarios I, II, and III for the rolling window approach.

### Resilience indicators

To measure resilience, we used metric-based resilience indicators based on critical slowing down (7, 10, 16). We specifically considered the performance of variance and autocorrelation as resilience indicators and performed significance tests using different methods for continuous monitoring and burst monitoring (see below).

#### Significance test for continuous monitoring – rolling window approach

For detrended continuous time series, we calculated the autocorrelation and variance with a rolling window in the classical way, with a window size of 50% of the time series, using an implementation in Matlab (https://git.wur.nl/sparcs/generic_ews-for-matlab). The trend in the indicators was evaluated using the Kendall tau value. We used a null model to assess the significance of the tau values. Specifically, we generated 100 surrogate data sets with the same power spectrum but randomized phases (17). The p-value was estimated as the fraction of the surrogate data sets where the tau value was higher than in the observed data. A trend was considered significant if its p-value was below the 5% threshold.

#### Significance test to compare bursts of data – bursts approach

If bursts of data were collected, we used the slope of a Theil-Sen regression of the indicators with the rank of each burst to assess a loss of resilience over time. Specifically, we calculated the indicators variance and autocorrelation for each burst and fitted a Theil Sen regression over the indicators of all the bursts. The slope of the regression allows to quantify a potential increase of the indicator over all the bursts. The Theil Sen regression is insensitive to outliers and is more robust than traditional regressions for skewed data. The slope is calculated as the median of the slopes of all pairs of points (18).

We used a sieve bootstrapping algorithm to assess the significance of the decrease in resilience in the burst approach, preserving the autocorrelation structure of the time series. The bootstrapping algorithm consists of two steps (19). After detrending the data, an AR(n) model is fitted, and the data residuals are determined. Second, the AR(n) model is used to generate new time series, where stochasticity is included by drawing residuals of the AR(n) model with replacement. We generated bootstrapped time series for each burst, and used these time series to calculate the value of the indicator. Then, we fitted a Theil-Sen regression over the indicators of the bootstrapped bursts and obtained the p-value of the probability of the slope being different from zero. A loss of resilience was considered significant if its p-value was below the 5% threshold.

#### Measure of the performance

To quantify the performance of both approaches, the analyses were repeated for 100 generated time series to estimate the true positive rate (sensitivity) and true negative rate (specificity). For the true positive rate, we used time series with an increasing c, representing a loss of resilience. We calculated the true positive rate as the proportion of repetitions resulting in a p-value below 0.05. For the true negative rate, we used time series with a fixed c = 0.5, representing stable time series. We defined the true negative rate as the proportion of repetitions resulting in a p-value above 0.05.

## Results

### Performance of the burst method vs the rolling window method

In our simulations, the burst approach signals an upcoming transition in more than 80% of the cases (Figure 2A) and produces false alarms in less than 15% of the cases (Figure 2B). Increasing the number of bursts reduces the true positive rate of variance (from 100% for 2 bursts to 92% for 8 bursts) and autocorrelation (from 100% for 2 bursts to 88% for 8 bursts) but does not seem to affect the true negative rate significantly. To compare the burst approach with the rolling window approach, we used three sampling scenarios for the rolling window approach (see methods, Table 1, Figure 2C). Truncating the time series (scenario I) hinders the performance of the rolling window approach for both indicators. On the other hand, reducing the resolution (scenario II) mainly affects the performance of autocorrelation for the rolling window approach. Thus, bursts can enhance the performance of autocorrelation by sampling with a high resolution without increasing the total number of samples. Not surprisingly, in the gold-standard scenario III, where neither effort nor time is limiting, the rolling window approach can signal upcoming transitions best, with a true positive rate of 100%.

**Figure 2.**
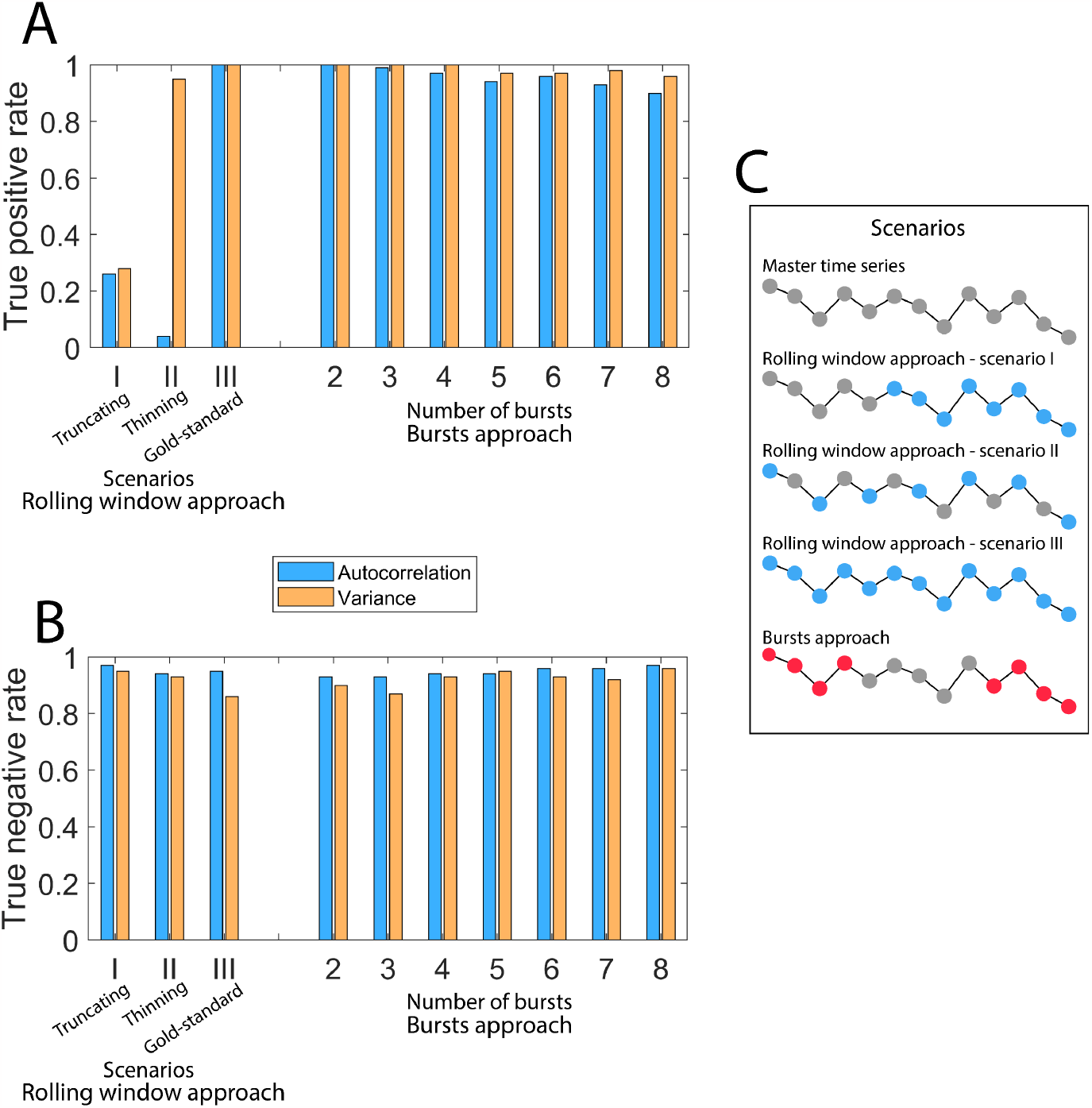
Performance of resilience indicators (autocorrelation and variance) depending on the sampling approach. For the rolling window approach, we use three scenarios for comparison (Table 1): scenario I, truncating, has the same total number of data points and resolution as the burst approach but covers a shorter gradient (i.e., only the last part of the master time series), scenario II, thinning, has the same number of data points and covers the full gradient from high to low resilience but a different resolution, and ‘gold-standard’ scenario III has the same resolution and gradient but a higher number of data points. For the burst approach, we vary the number of bursts. Except in scenario III of the rolling window approach, the total number of data points is L=1000 for both approaches, regardless of the number of bursts. Except in scenario II of the rolling window approach, the data points are subsampled with a daily resolution (Δ1=1 day). For the burst approach, the data points are equally allocated between the bursts, and the bursts are equally spaced in the time series. (A) True positive rate. The true positive rate is calculated in data simulated using the model (equation (1), parameter Table 1), with c increasing from 0.5 (high resilience) to 2.2 (low resilience) over 100 repetitions. (B) True negative rate. The true negative rate is calculated in simulated data with a fixed c=0.5 over 100 repetitions. (C) Illustration of the allocation of data points for the rolling window approach in scenarios I, II, and III and for the burst approach by subsampling from the master time series. Gray points are points in the master time series that are unused in that approach. Colored points are used.

### An empirical example: mood fluctuations before depression

We also tested the burst approach against a conventional rolling window approach using an existing dataset of mood changes (3). Using the rolling window approach on the full processed dataset, we found a significant rise in variance (p-value = 0.01) before the transition into depression, consistent with previous findings (3). Autocorrelation showed a weak signal when computed on the raw data, ignoring non-equidistant spacing (p-value = 0.056). However, significance increased when we used the daily mean value (p-value=0.03, see supplementary material), possibly due to the fact that this corrected for daily cycles and made data intervals equidistant. As aggregating per day left too little data to perform the burst analysis, we proceeded with variance to compare the bursts vs the moving window approach.

The results illustrate that, consistent with the findings from simulated data, the burst approach could signal an upcoming transition (Figure 3). When subsampling the data, the burst approach could detect the transition down to 160 data points for 2 bursts and 240 for 3 and 4 bursts. Thus, our real data example illustrates that, indeed, in this field where continuous monitoring can be too demanding, the burst approach can be a good alternative. Details on our methods and full results are presented in the supplementary material.

**Figure 3.**
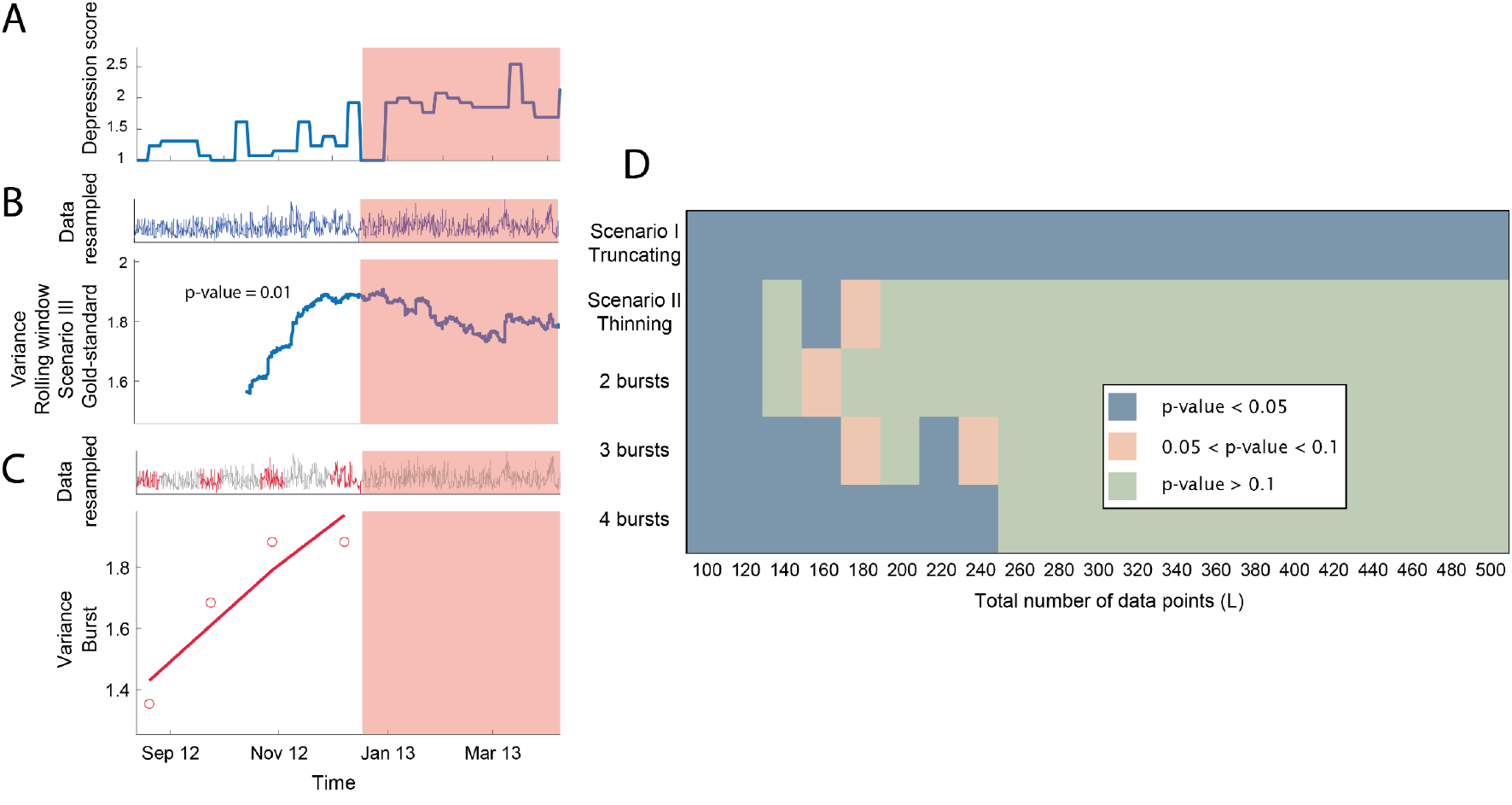
Analysis of the empirical dataset, mood fluctuations of a patient measured for 239 days through a 50-item questionnaire (A) Depression score over time, the shaded region indicates the tipping point. (B) Data used for the rolling window approach, scenario I, and trend of variance. The shaded region indicates the tipping point, 829 data points are available before the tipping point. (C) Data used for the burst approach and value of variance for each burst as well as the linear regression. (D) Comparison of the rolling window approach and burst approach in anticipating a depressive episode using mood data from (3) when subsampling from the master time series under scenario I: truncating, and II: thinning (rolling window approach) and for different number of bursts (burst approach). The performance of the approach is quantified using the p-value of the Ebisuzaki test (17) for the rolling window approach and of the slope being significantly different than zero for the burst approach. Scenarios for the rolling window approach are summarized in Table 1 and depicted Figure 2C.

### Effect of the sampling interval

We sampled from the simulated master time series to investigate the sampling interval (i.e., the interval between two samples within a burst) on the performance of the burst approach (Figure 4A and Figure 4B). Our results suggest that the sampling interval only affects the autocorrelation. To optimize the performance of autocorrelation, there is an optimal sampling interval. This is consistent with previous results (10) and additional analyses provided in the supplementary (Figure S2). Variance was hardly affected by the sampling interval.

**Figure 4.**
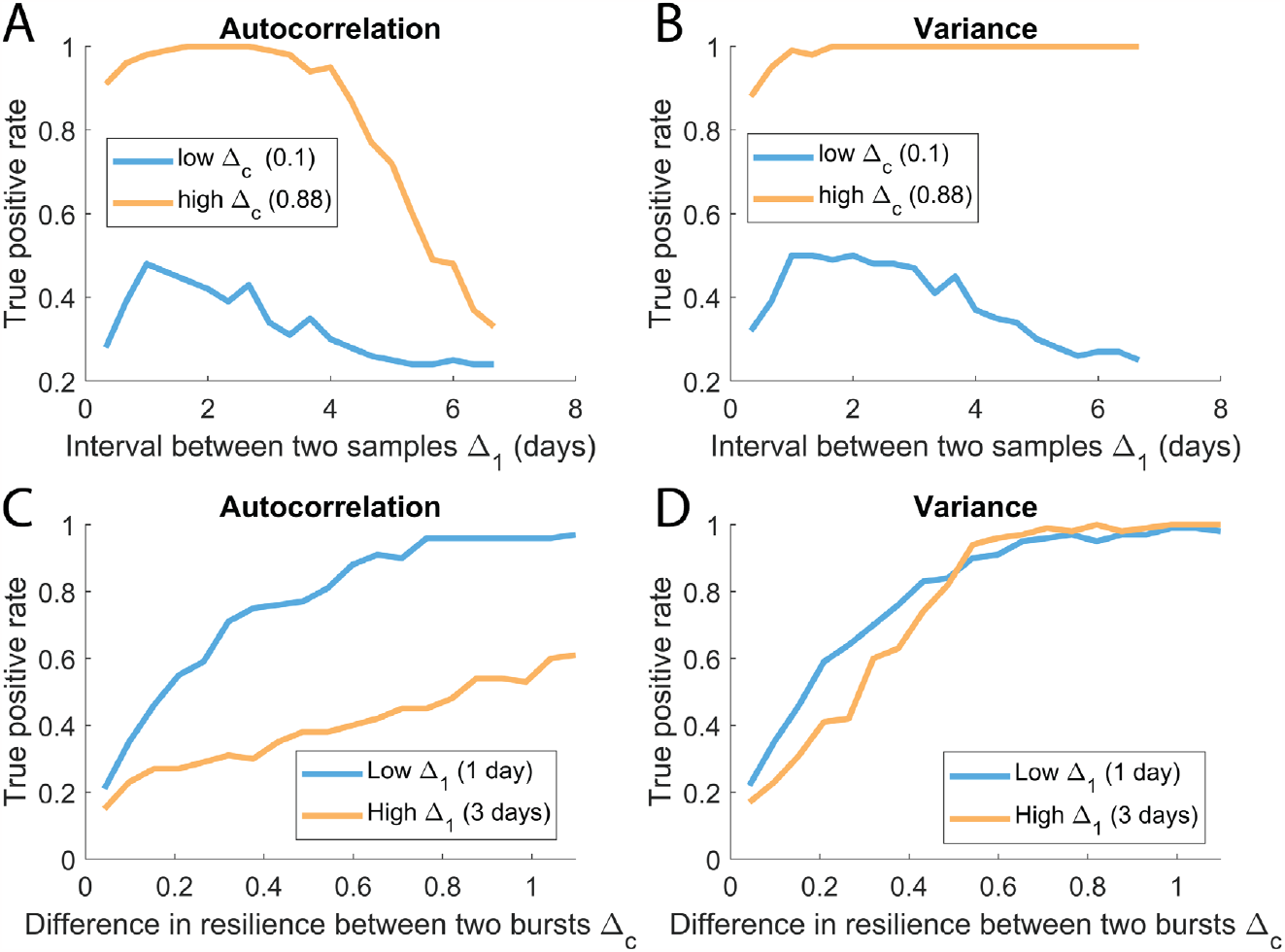
Effect of the sampling interval Δ1 (A and B) and the difference in resilience between two bursts Δc (C and D) on the performance of the bursts approach. The performance is quantified using the true positive rate over 100 repetitions, using subsamples from the master time series. The total number of data points was constant (L=1000) and spread equally between two bursts. (A) Performance of autocorrelation depending on the interval between two samples Δ1. We estimated that performance for two values of Δc. (B) Performance of variance depending on the interval between two samples Δ1. We estimated that performance for two values of Δc. (C) Performance of autocorrelation depending on the difference in resilience between two bursts Δc. We estimated that performance for two values of Δ1. (D) Performance of variance depending on the difference in resilience between two bursts Δc. We estimated that performance for two values of Δ1.

### Effect of the interval between bursts

Additionally, we investigated the effect of the interval between two bursts on the performance of the burst approach (Figure 4C and Figure 4D). In our model setup, a longer interval between two bursts implies a larger difference in resilience between the two bursts, which improved the detection of a loss of resilience for both variance and autocorrelation. Furthermore, there was no effect of the interaction of the sampling interval and the interval between two bursts, as the same patterns were observed for the different combinations of conditions.

### Effect of the number of bursts

To explore trade-offs in sampling strategies, we analyzed different combinations of choices under cost restraints.

#### Fixed total number of samples L

First, we investigated the case where the total number of samples L is fixed (Figure 5), as could be the case when each collected sample has a high cost. Under this constraint, a too-high number of bursts hampers the detection of a loss of resilience, especially for autocorrelation, as the bursts become too short and do not have enough data points to be representative of the state of the system. However, a lower number of bursts results in a later detection of the critical transition (Figure S7). Additionally, consistent with previous results (Figure 4A and Figure 4B), the performance of autocorrelation diminishes for a long sampling interval. These results suggest that there is an optimal way of allocating L samples when monitoring resilience in bursts, with high enough resolution and large enough bursts with a big enough interval between each burst. As more bursts increase the lead time of prediction but decrease the prediction performance, a compromise has to be found depending on the objectives of the study. Similar results were observed for different values of the total amount of data points *L* (Figure S5), where, not surprisingly, a decrease in the overall true positive rate was observed for a decrease in *L*.

**Figure 5.**
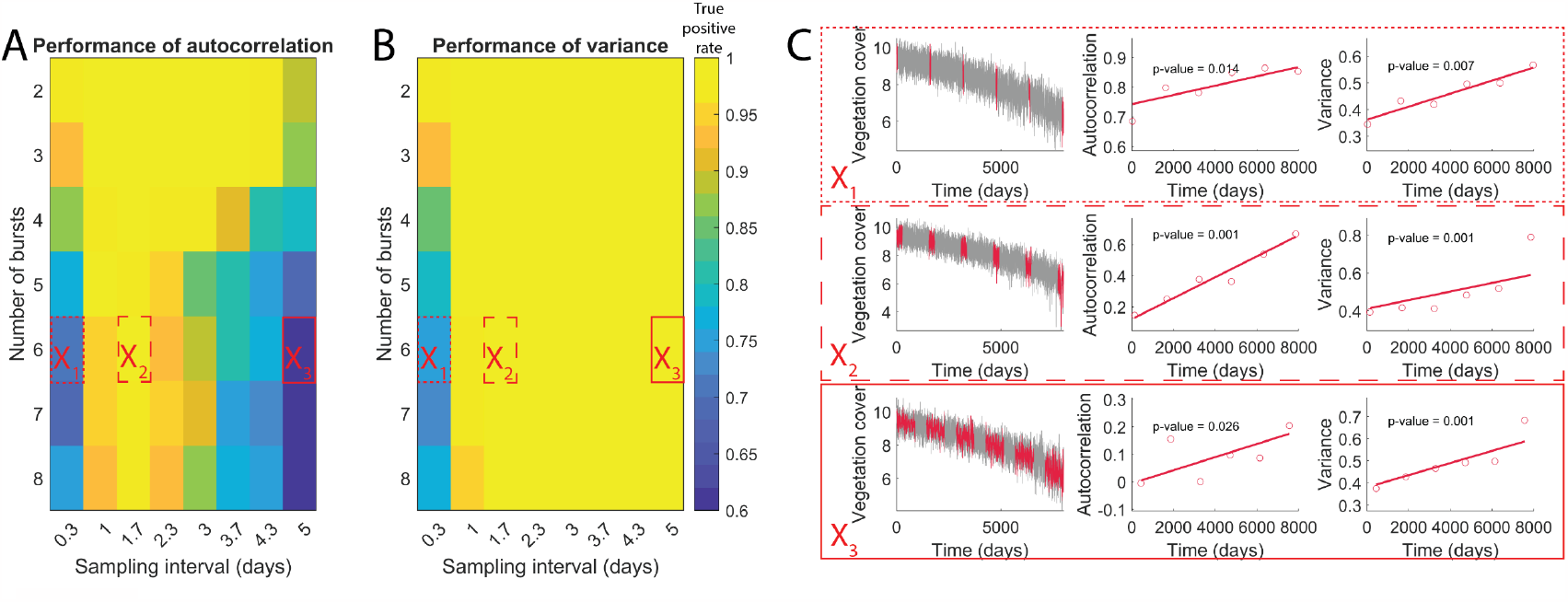
True positive rate of the bursts approach depending on the number of bursts n and sampling interval Δ1. The true positive rate is estimated using the time series generated with the vegetation model over 100 repetitions. The total number of data points was constant (L=1000), regardless of the number of bursts. The bursts are equally spaced in time. (A) Performance of the autocorrelation. (B) Performance of the variance. (C) Example of subsampling in the master time series for three combinations of parameters specified in A and B.

#### Fixed number of samples per burst λ

In addition, we investigated a scenario where costs are purely determined by the number of bursts, keeping the number of samples per burst constant (Figure 6). This scenario is relevant when an expensive device must be rented for the monitoring of a burst or a patient should come to the hospital for each burst. Opposed to the previous scenario, the number of bursts did not affect the performance, but still affected the lead time of prediction (Figure S8). However, under our assumption of a linear loss of resilience, large numbers of bursts coupled with large sampling intervals led to a shorter interval between the bursts and, thus, a smaller loss of resilience between each burst, hampering the signal. These results suggest that when the number of samples per burst is fixed, the sampling interval within the burst and the interval between two bursts are the only factors affecting the performance. Similar results were observed for different values of the amount of data points per burst *λ* (Figure S6), yet a decrease in the overall true positive rate was observed for a decrease in *λ*.

**Figure 6.**
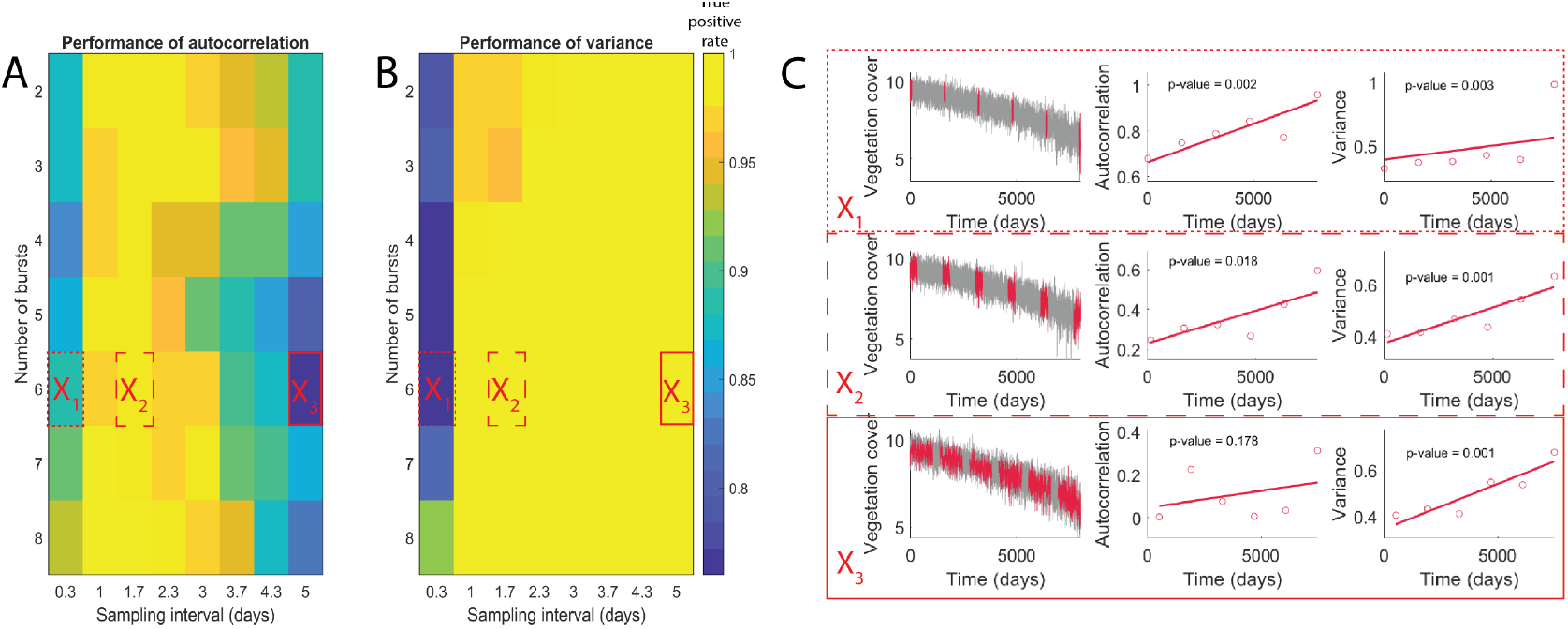
True positive rate of the bursts approach depending on the number of bursts n and sampling interval Δ1. The true positive rate is estimated using the time series generated with the vegetation model over 100 repetitions. The number of data points per burst was constant (λ=200), regardless of the number of bursts. The bursts are equally spaced in time. (A) Performance of the autocorrelation. (B) Performance of the variance. (C) Example of subsampling in the master time series for three combinations of parameters specified in A and B.

## Discussion

Our results imply that using bursts (i.e., two or more short high-resolution time series) to monitor resilience can be a powerful alternative to the more common sampling of long, continuous time series analyzed using a rolling window approach (Figure 1). This result was true for both simulated (Figure 2) and real-life data (Figure 3). Since reaching the data quality standards required for the rolling window approach can be fastidious (9, 20), the apparent power of the burst approach may expand opportunities to monitor resilience (Table 2).

**Table 2.** Example of applications for bursts and moving indicators of resilience.

|  | Objective | Monitoring in bursts | Continuous monitoring | Reference |
| --- | --- | --- | --- | --- |
| Vegetation | Anticipate a decline in vegetation due to overharvesting | Bursts of monitoring the fluctuations in vegetation cover over shorter periods of time (or spatial snapshots) | Continuous monitoring of the amount of vegetation | (27) |
| Depression | Anticipate episodes of depression using mood-tracking | Investigate a potential loss of resilience by using snapshots of mood fluctuations for a patient, reducing the dropout rate | Use continuous, long-term mood tracking and investigate the trend in resilience indicators | (2, 3) |
| Epidemiology | Anticipate epidemics using incidence time series | Compare incidence time series with previous years, or use bursts of active surveillance | Anticipate an outbreak in a specific area using long-term data | (5) |
| Species extinction | Anticipate population collapse using population abundance | Monitor population abundance during short time period with high resolution and compare with previous bursts | Monitor the population continuously during a long period | (4) |
| Resilience of human body: aging | Anticipate a decline in elderly using data from balance boards or grip strength for instance | Investigate a loss of resilience in a patient using bursts of data collected regularly by placing the patient on a balance board | In that case, it is impossible to monitor continuously the balance or grip strength of a patient | (22, 23) |
| Resilience of human body: chronic diseases | Anticipate acute episodes of chronic diseases using biomarker data over time | Collect bursts of biomarker concentration in a patient to investigate a loss of resilience | Estimate biomarker concentration regularly over a long time period (may cause inconvenience for patients and higher risk of dropouts) | (22, 28) |
| Resilience of human body: hypertension | Anticipate upcoming hypertensive crisis using blood pressure time series | Collect bursts of blood pressure in a patient | Estimate blood pressure regularly over a long time period (may cause inconvenience for patients and higher risk of dropouts) | (22, 29) |
| Resilience of human body: sports | Anticipate athletes' loss of resilience using physiological and psychological data | Collect bursts of physiological and psychological data to reduce the risk of dropout | Monitor physiological and psychological variables continuously over long time periods | (30) |
| Animal (livestock) resilience | Anticipate loss of resilience in livestock animals to increase animal welfare, using physiological variables | Collect bursts of physiological variables in livestock animals to reduce monitoring efforts by going to the farms less regularly for intense monitoring | Collect data regularly over long time periods to monitor a linear loss of resilience in the long run | (22, 31) |

Given a limited total number of data points, monitoring in bursts allows increasing the resolution of collected time series. High resolution is essential to detect changes in autocorrelation (Figure 4), which is a more direct and robust indicator of resilience than variance, which is affected by other factors such as environmental noise (21).

Bursts may, in particular, be useful for monitoring resilience of systems that are hard to measure continuously, such as patients. Monitoring resilience in human (mental) health can help anticipate critical transitions resulting in chronic diseases, depressive episodes, or loss of resilience due to unhealthy aging (22). Gijzel et al. (2019) used short time series collected with different individuals to show that autocorrelation and variance in postural balance time can discriminate between healthy and unhealthy aging in elderly. Although they compared time series from different individuals, their approach is comparable to our burst approach, where shorter time series are collected to compare resilience in a system over time.

We illustrated the potential value of a burst approach in a medical context, showing how a depressive episode could have been anticipated in a self-reported mood time series with much less burden to the subject than collecting the entire time series, in which a patient subject to depressive episodes monitored his daily life experiences up to ten times a day for 239 days in a row (Figure 3).

It is easy to think of other fields in which a burst approach might increase the power to detect changes in resilience that could precede critical transitions. For instance, in epidemiology, resilience indicators have been shown to signal upcoming disease outbreaks, but data requirements are hardly met, especially in periods of low transmission when the disease is not being monitored (24). Active surveillance, namely the active seeking of cases of a given disease in a population, provides ideal data to calculate resilience indicators. In active surveillance data, the prevalence of a disease is estimated with high accuracy. However, these data are extremely costly and not achievable in the long term. Bursts of active surveillance could allow to control if the risk of disease outbreak is increasing over time.

Also, key variables in ecological systems may be challenging and costly to monitor continuously. For instance, estimating species abundance in remote ecosystems requires repeatedly reaching the location to perform the measurements, as is the case for marine ecosystems where fish abundance time series may help estimate resilience (25). Indeed, Verbesselt et al. (2016) used bursts of data to compare the resilience of tropical forests at several locations. The authors showed that a higher autocorrelation was associated with lower resilience of a subsystem, consistent with our findings that the resilience of a system can be quantified by bursts of data in that system.

Although bursts of data can improve monitoring practice for a variety of systems, the power of the design will depend on trade-offs between the number of bursts, length of burst, sampling interval, and interval between two bursts (Figure 7). Optimal strategies depend on whether the costs per sample (Figure 5) or the costs per burst (Figure 6) determine the total sampling costs. In the first case, the number of bursts should still be limited to make sure that there are enough data points in each of the bursts. A large interval between two bursts, resulting in a large difference in resilience in the case of a linear loss of resilience, makes it easier to detect a difference between the bursts (Figure 4C and D). However, obviously, a large interval between subsequent bursts also increases the risk of missing the transition, as information about the resilience is not updated between the bursts. To help overcome that challenge, one could determine the interval between the bursts adaptively: starting with bursts close to one another and then increasing the interval between two bursts if there was little change in resilience visible.

**Figure 7.**
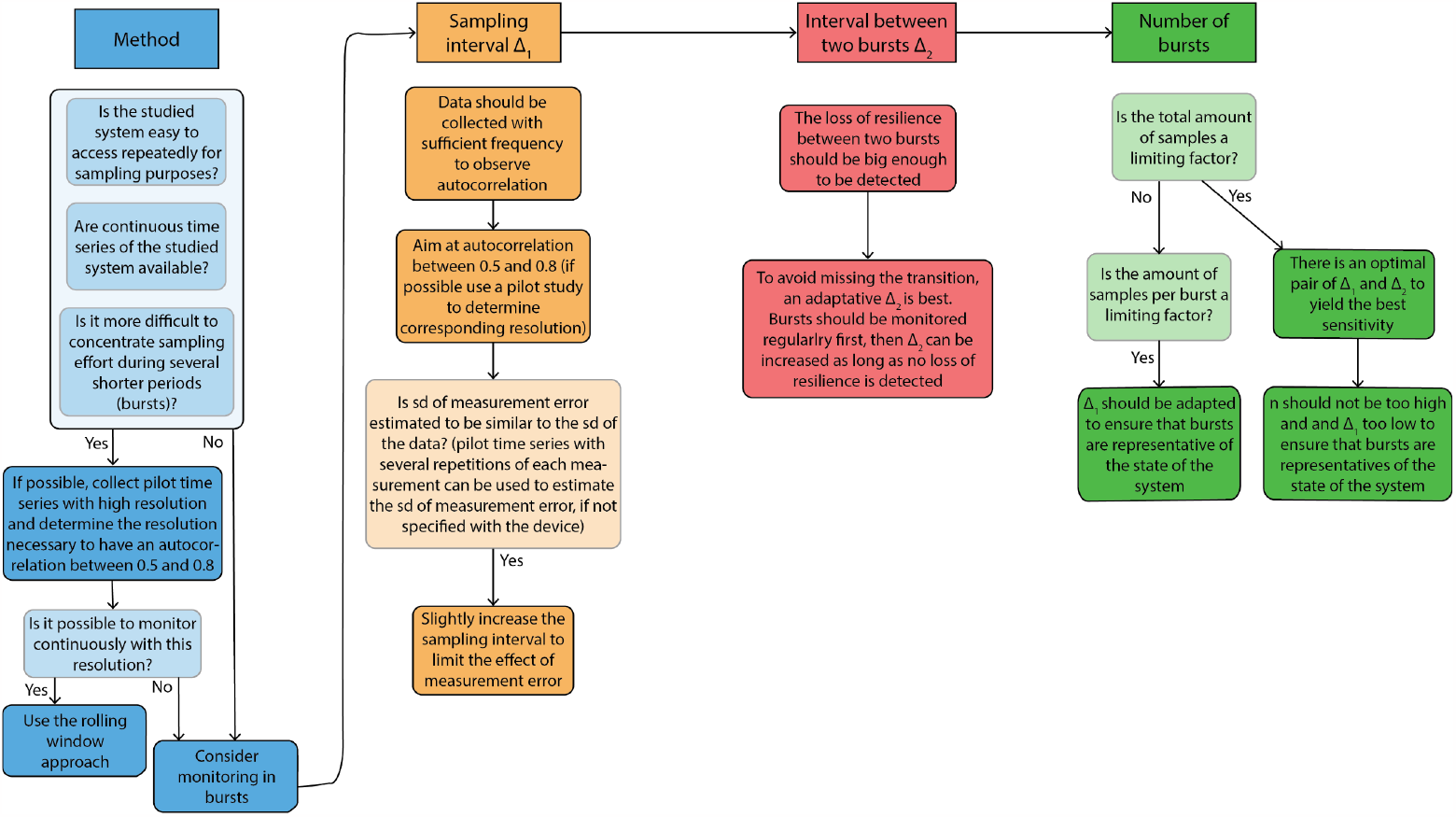
Decision tree to allocate sampling resources when monitoring a potential loss of resilience, when sampling resources are limiting.

In summary, while the power and optimal design of the approach depend on the context, repeated bursts typically outperform continuous monitoring of resilience. The exception being situations where there are no cost or practical limitations to data collection.

## Supporting information

All supplementary material, text, figures S1 to S8 and Table S1

## Data Availability

All relevant data can be found at https://github.com/cla-delec/Monitoring_resilience_bursts and software at https://git.wur.nl/sparcs/generic_ews-for-matlab

