## Supplementary material for "Monitoring Resilience in Bursts": All supplementary material, text, figures S1 to S8 and Table S1

### Stability analysis of the model

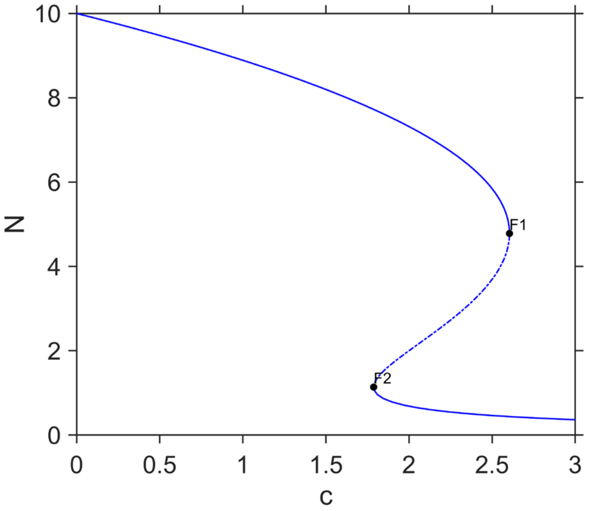


*Figure S1 Bifurcation of the deterministic version of the model with varying parameter c. This model has one stable high vegetation state for low levels of c, two stable states for intermediate values of c, and a low vegetation state for high c. At F1 (c=2.6) and F2 (c=1.8), the system crosses a fold bifurcation.*

### Effect of measurement error on the autocorrelation

*Autoregressive model*

We used an autoregressive model to investigate the effect of the sampling interval $\Delta_1$ and the amplitude of the measurement error $\sigma_{ME}$ on the accuracy of the estimation of autocorrelation.

$$n(t+1) = \alpha(n(t) - n_0) + n_0 + \sigma\varepsilon$$

(2)

In the model, α is the autocorrelation with lag 1, namely the correlation between two time points delayed with a lag of one, $n_0$ represents the initial condition and $\sigma\varepsilon$ is white noise to reproduce intrinsic stochasticity. In this model, each datapoint depends on the previous datapoint. This model is a direct representation of the autocorrelation and helped us studying the effect of other factors on the autocorrelation straightforwardly. After simulating the time series, we added measurement error by sampling from a standard normal distribution for each time step, multiplied by an amplitude factor $\sigma_{ME}$. We used α=0.99, $n_0$=900 and $\sigma\varepsilon = 1$, and several values for $\sigma_{ME}$=0, 1, 3, 5.

The effect of the sampling interval $\Delta_1$ and the amplitude of measurement error $\sigma_{ME}$ on was then investigated (Figure S1). Without measurement error, the estimation of autocorrelation is more accurate for a short sampling interval. However, including measurement error resulted in an optimal sampling interval to measure the autocorrelation with the best accuracy.

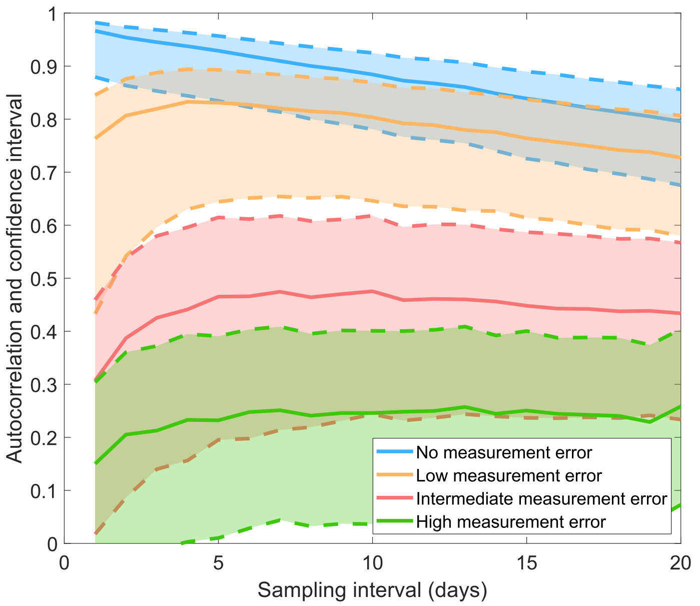


*Figure S2 Autocorrelation and its confidence interval depending on the sampling interval and measurement error, calculated in time series generated with the autoregressive model*

### Anticipation of a depressive episode in a dataset

*Trend in autocorrelation*

Using the dataset described in Methods – Dataset, we tried to anticipate a depressive episode using time series of daily experiences of a patient. Autocorrelation showed a weak signal due to the inconstant sampling interval, as the daily experiences were sampled at random times of the day and no monitoring was done at night. To overcome this issue, we tried different pre-processing of the data: detrend for daily cycles (by subtracting the mean of each day), not take into consideration in the calculations consecutive data points measured with an interval bigger than 4 hours and 6 hours, aggregate the data per day (using mean), reducing the resolution (scenario B for the rolling window approach, see methods) and use all the data. The strongest trend in autocorrelation was observed when using aggregated data per day, suggesting that indeed daily cycles hamper the use of autocorrelation for this dataset.

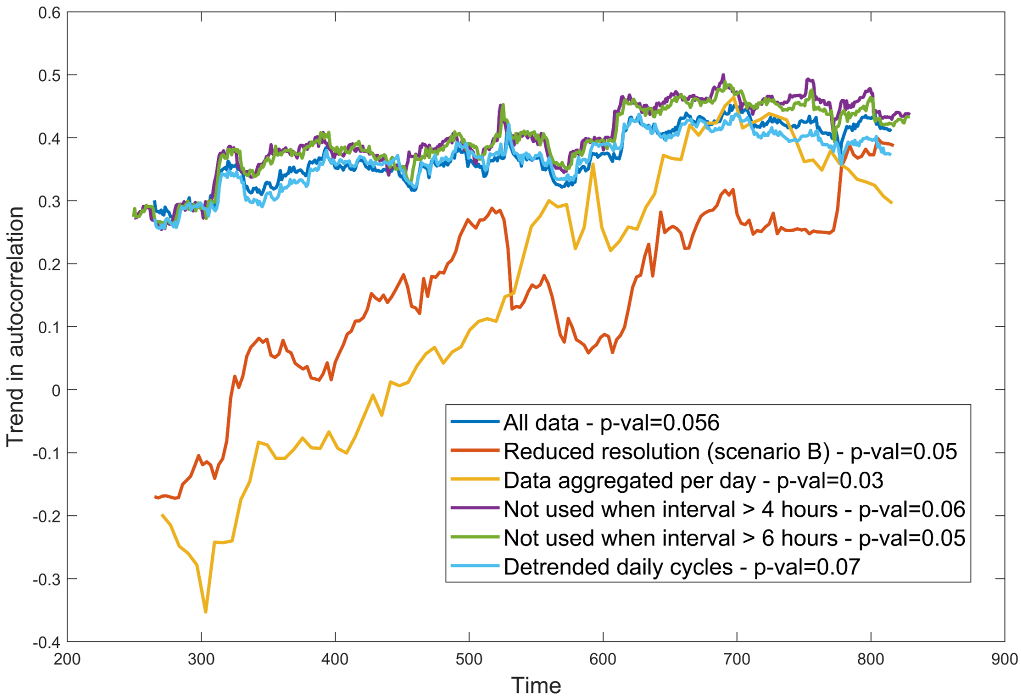


*Figure S3: Estimation of the trend in autocorrelation for different pre-processing of the data: (1) dark blue: raw data, (2) orange: scenario II of the rolling window approach (see methods), resolution is reduced by subsampling every 3$^{rd}$ datapoint, (3) yellow: aggregate data per day using the mean, (4) purple: consecutive data points measured with an interval bigger than 4 hours are not taken into consideration in the calculation of autocorrelation, (6) green: purple: consecutive data points measured with an interval bigger than 6 hours are not taken into consideration in the calculation of autocorrelation, (7) light blue: data are detrended for daily cycles by subtracting the mean of each hour to the corresponding data points.*

### *Sensitivity to the size of the rolling window*

We investigated the sensitivity of the trend in autocorrelation and variance to the size of the rolling window. Variance was insensitive to the size of the rolling window, but autocorrelation was. We picked a rolling window of 30% for all analyses.

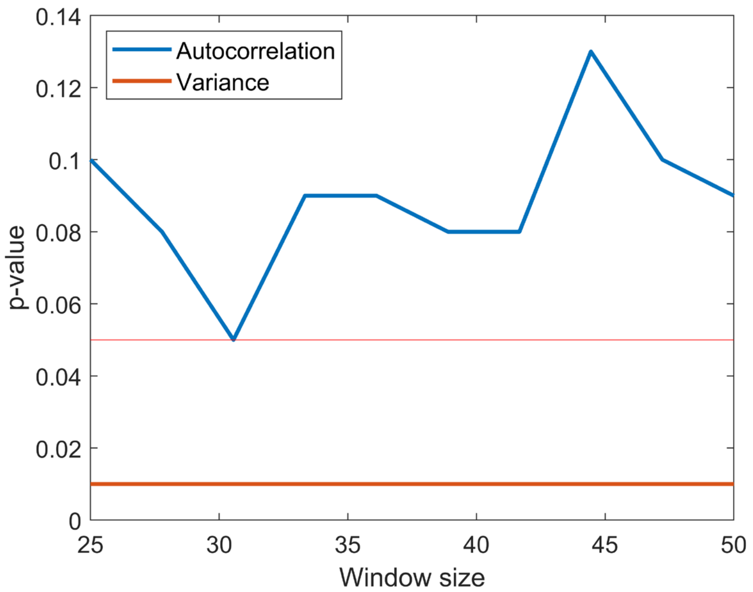


*Figure S4: p-value of the trend of autocorrelation and variance for different sizes of the rolling window*

*Scenarios*

Additionally, we investigated several tangible monitoring scenarios to illustrate how bursts of measurement could have reduced the sampling intensity for the patient.

Reducing the sampling to twice two weeks of measurements would have still signaled an upcoming transition, and would have been more convenient for the patient.

*Table S1: results of different sampling scenarios to anticipate depressive episodes in the daily experiences dataset*

| Number of bursts | Duration of one burst | Total number of datapoints | p-value of variance |
|---|---|---|---|
| 2 | 2 weeks | 158 | 0.034 |
| 2 | 12 days | 135 | 0.043 |
| 3 | 2 weeks | 259 | 0.02 |
| 4 | 2 weeks | 356 | 0.009 |

### Effect of the total number of data points on the optimal sampling strategy when the total number of data points is fixed

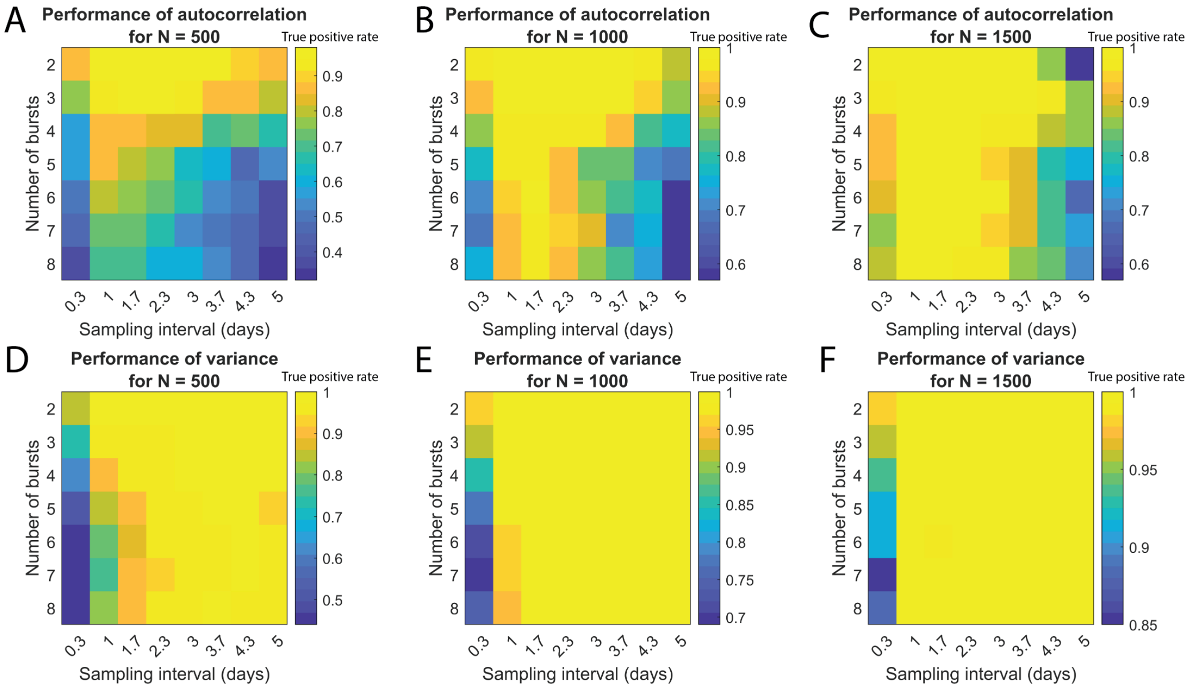


*Figure S5 True positive rate of the bursts approach depending on the number of bursts n and sampling interval Δ1, for three values of total data length L (500, 1000 and 1500). The true positive rate is estimated using the time series generated with the vegetation model, over 100 repetitions. The total number of data points was constant regardless of the number of bursts. The bursts are equally spaced in time, from the start to the end of the time series. (A) Performance of the autocorrelation. (B) Performance of the variance.*

### Effect of the number of data points per burst on the optimal sampling strategy when the number of data points per burst is fixed

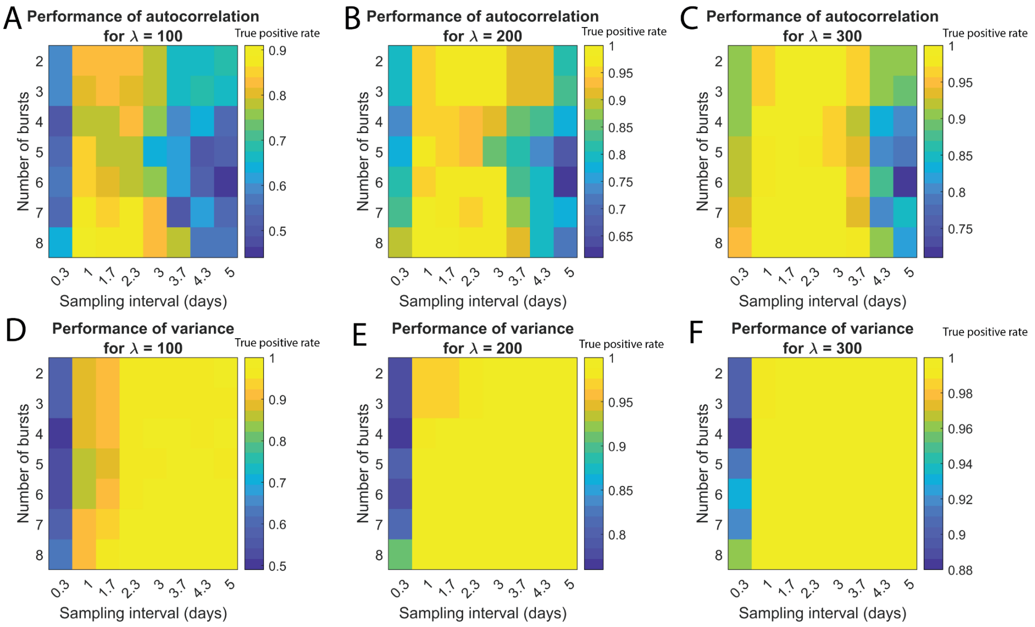


*Figure S6 True positive rate of the bursts approach depending on the number of bursts n and sampling interval $\Delta 1$, for three values of number of values per burst $\lambda$ (100, 200 and 300). The true positive rate is estimated using the time series generated with the vegetation model, over 100 repetitions. The number of data points per burst was constant regardless of the number of bursts. The bursts are equally spaced in time, from the start to the end of the time series. (A) Performance of the autocorrelation. (B) Performance of the variance.*

### Lead time of prediction

Due to the distribution of bursts in the master time series, a low number of bursts showed a greater performance. This is partly explained by the fact that bursts are spread from the start to the end of the master time series. Thus, in our analyses, a lower amount of bursts results in a larger interval between the bursts. However, in real-life, one may want to detect an upcoming critical transition as early as possible. Then, every time a new burst of data is obtained, the analysis of resilience indicators will be run again with this new data. In that case, having more bursts can have the advantage of being able to anticipate the upcoming transition earlier. We estimate the lead time by measuring the minimal number of bursts within a subset still leading to the detection of a significant loss of resilience, and then measure the corresponding distance between the last data point and the critical transition. We measure the lead time over 100 repetitions and plot the average earliness over all the repetitions (figure S6 and S7). By definition, the lead time is zero for two bursts because we sample bursts from start to end of the time series, thus the second burst is placed right before the critical transition resulting in a lead time of 0 days. We observe that a higher number of bursts leads indeed to an earlier detection of the critical transition, highlighting an advantage of monitoring more bursts.

*When the total number of data points L is fixed*

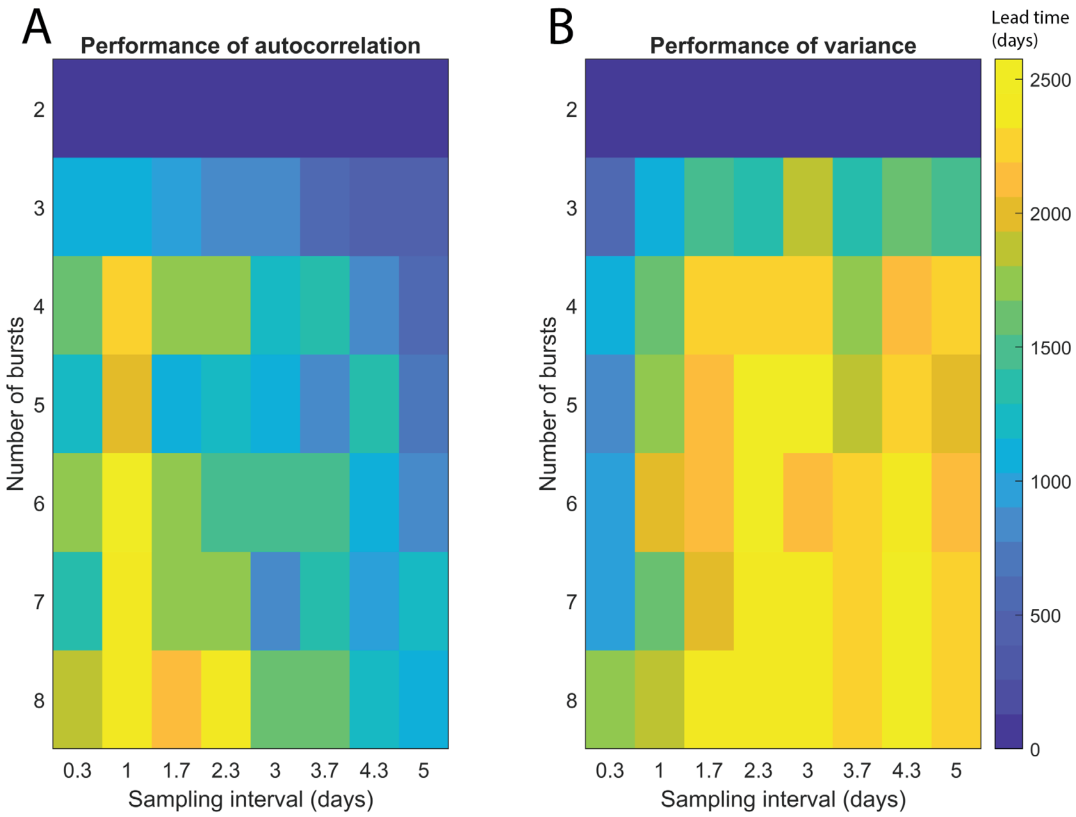


*<u>Figure S7</u> Lead time of prediction depending on the number of bursts n and sampling interval Δ1. The lead time is defined as the time between the last collected data points leading to a significant detection of the loss of resilience and the critical transition. We used the time series generated with the mode for the analyses, over 100 repetitions. The total number of data points was constant (L=1000), regardless of the number of bursts. The bursts are equally spaced in time. (A) Performance of the autocorrelation. (B) Performance of the variance.*

*When the number of data points per burst λ is fixed*

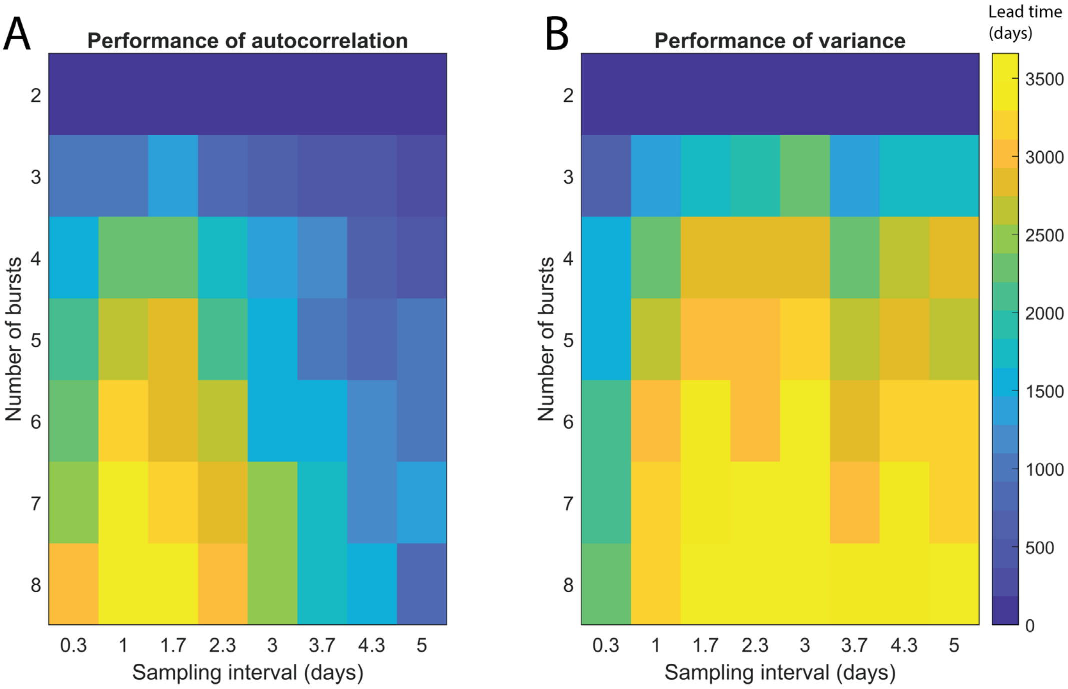


*Figure S8 Lead time of prediction for autocorrelation and variance depending on the number of bursts n and sampling interval Δ1. The lead time is defined as the time between the last collected data points leading to a significant detection of the loss of resilience and the critical transition. We used the time series generated with the mode for the analyses, over 100 repetitions. The number of data points per burst was constant (λ=200), regardless of the number of bursts. The bursts are equally spaced in time. (A) Performance of the autocorrelation. (B) Performance of the variance.*
